# High-force magnetic tweezers with hysteresis-free force feedback

**DOI:** 10.1101/862342

**Authors:** D. Kah, C. Dürrbeck, W. Schneider, B. Fabry, R. C. Gerum

## Abstract

Magnetic tweezers based on solenoids with iron alloy cores are widely used to apply large forces (~100 nN) onto micron-sized (~5 *μ*m) superparamagnetic particles for mechanical manipulation or microrheological measurements at the cellular and molecular level. The precision of magnetic tweezers, however, is limited by the magnetic hysteresis of the core material, especially for time-varying force protocols. Here, we eliminate magnetic hysteresis by a feedback control of the magnetic induction, which we measure with a Hall sensor mounted to the distal end of the solenoid core. We find that the generated force depends on the induction according to a power-law relationship, and on the bead-tip distance according to a stretched exponential relationship. Together, both relationships allow for an accurate force calibration and precise force feedback with only 3 calibration parameters. We apply our method to measure the force-dependence of the viscoelastic and plastic properties of fibroblasts using a protocol with stepwise increasing and decreasing forces. We find that soft cells show an increasing stiffness but decreasing plasticity at higher forces, indicating a pronounced stress stiffening of the cytoskeleton. By contrast, stiff cells show no stress stiffening but an increasing plasticity at higher forces. These findings indicate profound differences between soft and stiff cells regarding their protection mechanisms against external mechanical stress. In summary, our method increases the precision, simplifies the handling and extends the applicability of magnetic tweezers.

**SIGNIFICANCE:** Magnetic tweezers are widely used, versatile tools to investigate the mechanical behavior of cells or to measure the strength of receptor-ligand bonds. A limitation of existing high-force magnetic tweezer setups, however, is caused by the magnetic hysteresis of the tweezer core material. This magnetic hysteresis requires that the tweezer core must be de-magnetized (de-Gaussed) prior to each measurement, and that flexible force protocols with decreasing forces are not possible. We describe how these limitations can be overcome with a force feedback though direct magnetic field measurement. We demonstrate the applicability of our setup by investigating the visco-elastic and plastic deformations of fibroblasts to forces of different amplitudes.

## INTRODUCTION

Magnetic tweezers are a widely used tool for manipulation, force measurements and force application at the cellular and molecular level (1). Magnetic tweezers have been applied for cell rheology measurements (2), for investigating the binding strengths of specific membrane proteins (3) or for manipulating individual DNA molecules (4). Their force range is typically on the order of 10^−3^–10^4^ pN (5), surpassing optical tweezers (6) and dielectrophoresis-based tweezers (7) by several orders of magnitude.

The simplest design for magnetic tweezers is an electromagnet in the form of a solenoid with a core made out of a material with high magnetic permeability, such as soft iron. The core is tapered on one side to a sharp tip with a radius of typically less than 10 *μ*m. At the tip, a high gradient magnetic field is formed, which attracts nearby superparamagnetic beads (8). A major drawback, however, has been the effect of magnetic hysteresis of the core material (9).

Magnetic hysteresis implies that the magnetization of the core material does not only depend on the present solenoid current but also on its history. Practically, this means that the relationship between solenoid current and the generated force is difficult to predict and is typically calibrated only for a single, specific current protocol, e.g. for increasing currents after core de-magnetization (de-Gaussing). Thus, after each measurement, the remanent magnetization of the solenoid core must be eliminated by applying a sinusoidally alternating solenoid current that is reduced in amplitude over time (8, 10). However, even when performed very quickly, de-Gaussing always causes a large, sudden, distance-dependent force application to nearby beads. Alternatively, a smaller current in opposite direction can be applied to compensate for the remanent magnetic field of the core, but the magnitude of this counter-current needs to be specifically calibrated for each force protocol (11).

Another common strategy to mitigate the problems associated with magnetic hysteresis is to use a core made out of Mu-metal, an 80% nickel-iron-molybdenum alloy optimized for low hysteresis. However, Mu-metal has a considerably lower maximum magnetic induction compared to conventional iron alloys (12, 13) and can regain hysteretic behavior after machining, e.g. after sharpening the tip with a grinder.

It is also possible to eliminate the high-permeability core altogether and to generate the magnetic field and field gradient by a pair of coaxial coils with opposite polarity (14). In contrast to conventional magnetic tweezers, this system greatly simplifies the control of the applied magnetic force, which can be calculated by an analytical relationship that only depends on the coil current. Nevertheless, this system is unsuitable for many biophysical applications due to its low maximum force of approximately 2 pN for 4.5 *μ*m beads.

Finally, the remanent magnetic field of the core material can be compensated by a feedback circuit that regulates the magnetic induction of the core material instead of the magnetic current (15). The magnetic induction is measured with a Hall probe, and the magnetic current is then adjusted until the desired magnetic induction is reached. Such a system has been previously employed in a 4-pole tweezer setup where the primary function of the induction feedback was to reduce the magnetic cross-talk among the solenoids but not to compensate for the hysteresis of the core material. The maximum force of this setup was approximately 1 pN for 2.8 μm beads, which is too low for many applications.

In this study, we present a single-pole magnetic tweezer system with Hall probe-based induction feedback that is able to achieve much higher magnetic forces. By compensating the magnetic hysteresis of the core material, we can apply arbitrary force protocols with an amplitude of up to 100 nN to 5 *μ*m superparamagnetic beads. We also describe an empirical equation with only 3 free parameters that captures the distance/force relationship as a function of the magnetic induction for different superparamagnetic beads and for different core materials. This equation can be incorporated into the feedback loop to perform controlled force experiments. To illustrate the applicability of the technique as a tool in cell biological studies, we applied increasing and decreasing force steps on murine embryonic fibroblasts for evaluating the non-linear (force-dependent) viscoelastic and plastic cell rheology.

## MATERIALS AND METHODS

### System design

The core of the solenoid is a 4.5 mm diameter, 100 mm long cylinder made either out of St37 steel or Mu-metal (Vacuumschmelze GmbH, Hanau, Germany) (Fig. 1a-c). One side of the cylinder is tapered to a sharp tip (opening angle 60°, tip radius < 5 *μ*m) by grinding with a precision drill grinder while the cylinder is continuously turned by a milling machine (Fig. 1c inset) (for details see (16)). In the following, we refer to the sharp-tipped solenoid core as the needle.

**Figure 1:**
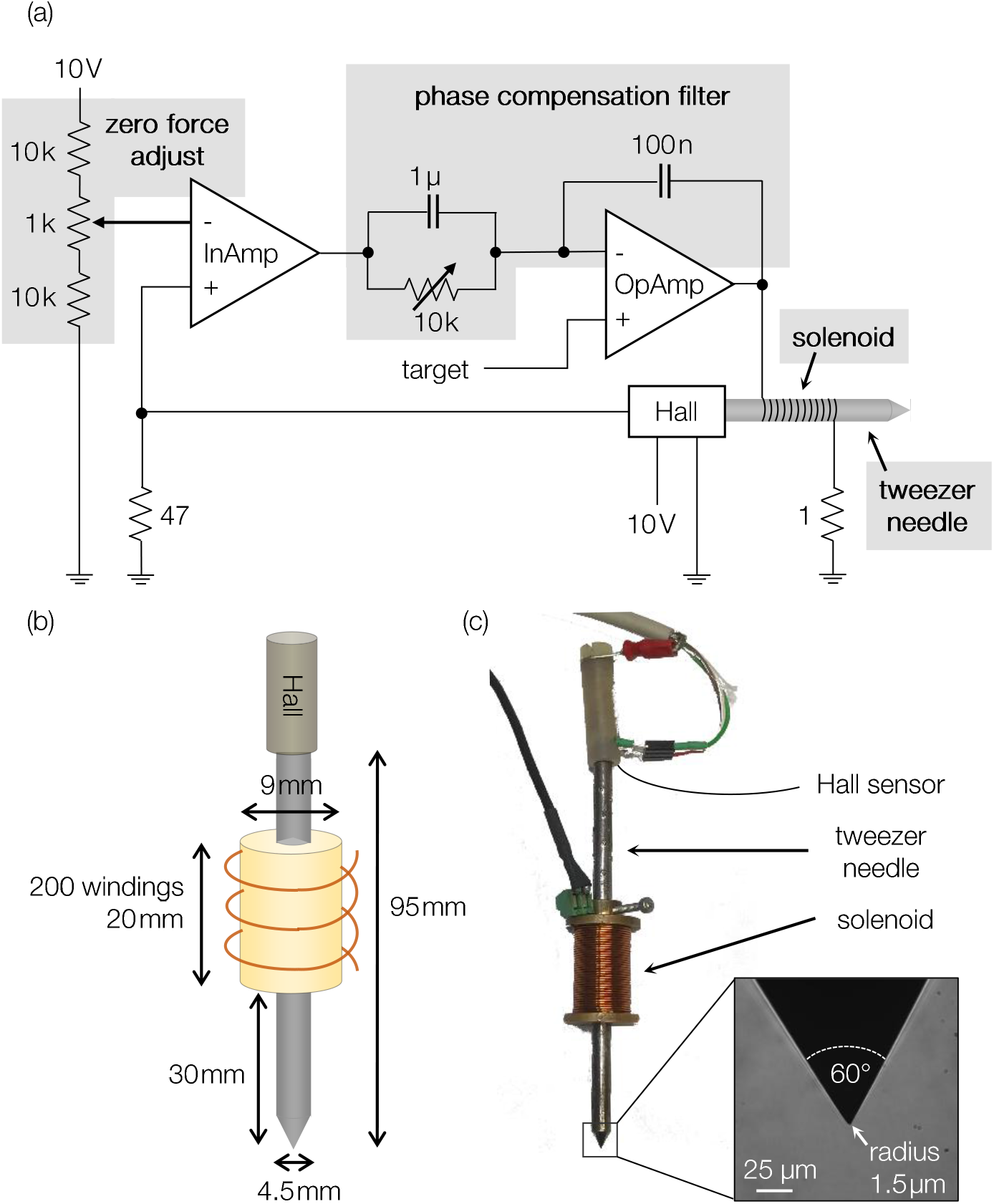
Electronic circuit and hardware. (a) The feedback control loop consists of three main components: a Hall probe to continuously measure the magnetic induction, an instrumentation amplifier to process the Hall signal, and a high-voltage/high-current operational amplifier to control the solenoid current. The operational amplifier compares the target value to the value measured by the Hall sensor. Through the feedback loop, the operational amplifier adjusts the solenoid current until the measured value from the Hall probe equals the target value. Zero force can be adjusted with a potentiometer. (b-c) The magnetic tweezer needle is magnetized by a solenoid (200 windings). The generated magnetic field is sensed by a Hall probe that is mounted onto the rear end of the magnetic tweezer needle. The needle has a sharp tip with an opening angle of 60° and a radius of 1.5 *μ*m to generate a high magnetic field gradient.

A Hall sensor (SS495A, Honeywell, Charlotte, NC) for measuring the magnetic induction was mounted at the rear (blunt) end of the needle. The needle is inserted in a solenoid with ~200 windings (24 gauge copper wire) on a brass body. The solenoid current is supplied by a high-current operational amplifier (OPA549T, Texas Instruments, Dallas, TX).

The needle is attached to a micromanipulator (InjectMan NI2, Eppendorf, Hamburg, Germany) to allow for precise movements of the needle tip. The position of the needle tip relative to a magnetic bead (for example a bead that is attached to a cell or dispersed in oil) is measured with an inverted bright field microscope equipped with a 40x 0.6 NA long working distance objective (Leica, Wetzlar, Germany). A specially manufactured non-magnetic objective is used because conventional lenses with nickel/chrome housings can distort the magnetic field in the object plane (17). Images are acquired with a CCD camera (Orca-Spark, Hamamatsu Photonics, Hamamatsu, Japan), which is triggered by a data acquisition card (NI-6052E, National Instruments, Asutin, TX), typically at a rate of 50 frames/sec. The analog output of the data acquisition card provides the set-point of the magnetic induction for the feedback loop circuit described below. This way, image acquisition can be synchronized to the force protocol, which is particularly important when recording fast movements of beads after sudden force changes.

To achieve a high frame rate, only a region of interest of 1920×128 pixels (corresponding to 310 x 21 *μ*m) is recorded during the measurement, which is sufficient to extract the bead-tip distance and the bead trajectory. Images are transferred to a computer via a USB 3.0 connection and evaluated with a custom software written in Python (18) using the Micromanager framework (19). Bead tracking is performed using an intensity-weighted center-of-mass algorithm (20). The position of the needle tip is determined from the image by thresholding, followed by erosion and dilation operations, yielding a binary image of needle and background. For every acquired frame, the bead position, the bead-to-tip distance, and the solenoid current are stored in an SQLite database file.

### Design of the control loop

The centerpiece of the control loop is a high-current operational amplifier operated as a non-inverted amplifier. The target voltage of the amplifier (at the non-inverted input) is provided by a data acquisition card. The feedback voltage (at the inverted input of the amplifier) is the amplified and phase-compensated magnetic induction signal measured by the Hall sensor. The amplifier output is connected to the solenoid. Positive feedback from high-frequency sources is suppressed with a 100 nF capacitor connected between the output and the inverted input of the amplifier (Fig. 1a).

The Hall sensor is a ratiometric linear sensor operated with a stabilized input voltage of 10 V supplied by a voltage reference (AD 587, Analog Devices), and has a measuring range of −67 to 67 mT. At zero magnetic field, the output voltage of the sensor is half the operating voltage, i.e. 5 V. This offset is removed with a resistor network in combination with an instrumentation amplifier (INA114AP, Texas Instruments, Dallas, TX) with unity gain.

The 1 kΩ potentiometer of the resistor network is used to tune the zero force point for a zero induction input voltage as follows: superparamagnetic beads with a radius of 5.09 *μ*m (microParticles GmbH, Berlin, Germany) are dispersed in water at a concentration of 5 ‧ 10^6^ particles/mL, and the magnetic tweezer needle is briefly magnetized to attract beads. The potentiometer of the resistor network is then adjusted until the beads detach from the needle tip when the microscope stage is abruptly moved.

Because the Hall probe has a response time of 3 ms, a phase-compensation RC filter is needed to suppress positive feedback, which can give rise to high-frequency (> 1 kHz) current oscillations. The 10 kΩ potentiometer of the RC filter is tuned to minimize response time, current overshoot and current oscillations when a square wave signal (1 V, 10 Hz) is applied as the target signal (Fig. 2c).

**Figure 2:**
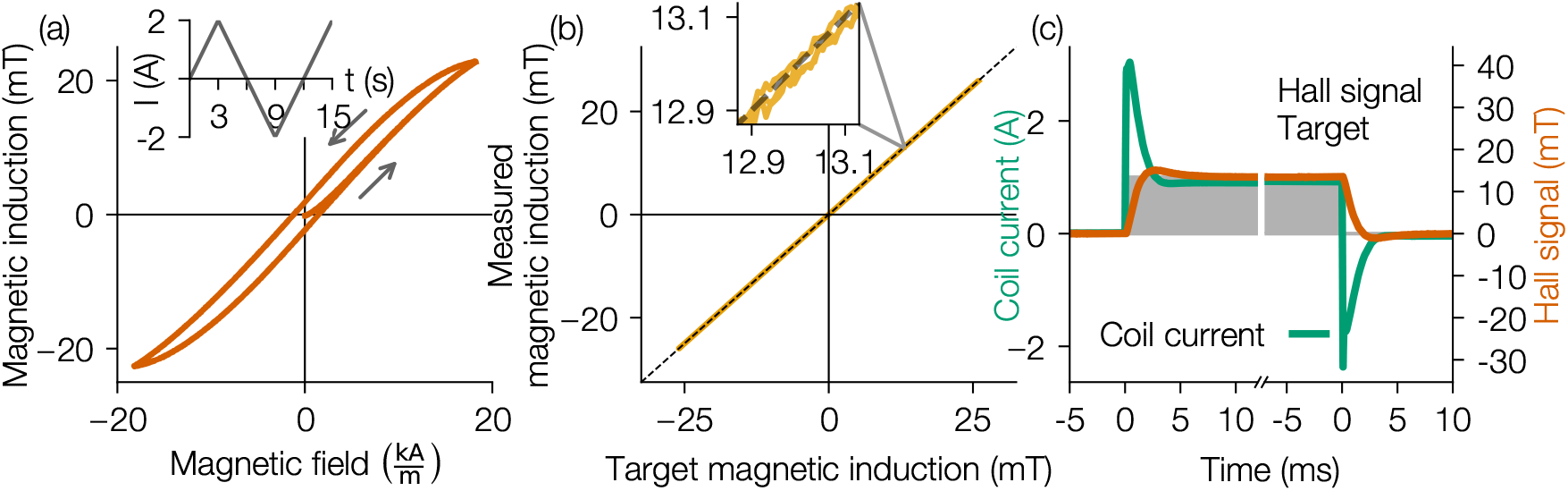
Hysteresis compensation and high-frequency response of the control loop. (a) Magnetization curve of a ST37 tweezer needle measured with a Hall probe in response to a triangular current protocol (current protocol shown in inset). (b) Magnetic induction measured with a Hall probe versus target induction for a triangular protocol when the induction feedback loop is turned on. Dashed line shows the line of identity. Inset shows the noise level of 18 *μ*T. (c) Step response of the control loop (induction measured with Hall probe (orange) and solenoid current (green)) for a square wave target signal (gray).

### Validation of hysteresis compensation

To characterize the low-frequency response of the control loop, we first recorded the hysteresis of the needle material (St37 steel) with the magnetic feedback turned off. For this purpose, we applied a triangular current protocol (period time 12 s, amplitude 2 A, Fig. 2a inset) and measured the magnetic induction with a Hall probe (Fig. 2a). As expected, we found a non-linear relationship between the magnetic field and the magnetic induction, and a pronounced hysteresis (coercivity of 1.5 mT). With the magnetic feedback turned on, we found a linear response free of hysteresis, with a deviation (rms) between target and measured magnetic induction of 18 *μ*T that remained constant over the entire range of input currents (Fig. 2b).

To characterize the high-frequency response of the control loop (see Fig. 2c), we recorded the coil current and the Hall signal in response to a square wave target signal (1 V, 10 Hz). The induction measured with the Hall probe approached the target value within 3 ms with a small overshoot (<10%) but without noticeable oscillations. The coil current displayed a larger overshoot during the first 3 ms after a sudden change in the target induction, which reflects the energy needed to overcome the hysteretic behavior of the needle and to some degree is also a consequence of the finite response time of the Hall probe.

### Force Calibration

The force that is exerted by the magnetic tweezers on a bead depends on the bead properties (e.g. bead radius, susceptibility and volume fraction of magnetic nanoparticles), and the needle properties (e.g. susceptibility of the core material and tip geometry). For a given needle-bead-combination, the force exerted on the bead moreover depends on the solenoid current and the distance between bead and needle tip (11). Below, we describe a calibration procedure that allows the user to apply controlled forces.

Beads were dispersed in a viscous liquid of known viscosity. Here, we used poly-dimethylsiloxane (PDMS) oil (Sigma-Aldrich, St. Louis, MO) with a dynamic viscosity of 9.65 or 28.95 Pa·s. We compared two types of superparamagnetic beads with different iron contents and functionalizations, namely 1) epoxylated beads with a diameter of 4.5 *μ*m and 20% iron oxide content (Dynabeads M-450, Invitrogen AG, Carlsbad, CA), and 2) carboxylated beads with a diameter of 5.09 *μ*m and 80% iron oxide content (microParticles GmbH, Berlin, Germany). In the following, these two types of beads are abbreviated as Fe20 and Fe80, respectively. For a calibration measurement, the beads dissolved in PDMS were filled into a glass dish (cf. Supporting Material S1 for a detailed experimental protocol) and moved with a manual microscope stage in such a way that a single bead was positioned 50 *μ*m in front of the needle tip. We then applied a constant magnetic induction for 1 s, followed by a zero induction pause for 1 s, and repeated this on-off-sequence until the bead reached the tip. Then another bead was selected, positioned 50 *μ*m in front of the needle tip, and the procedure was repeated with a different magnetic induction amplitude between 1.3 mT and 20.8 mT.

In response to the square wave pattern of the magnetic induction, the beads performed a stop-and-go-like motion. Bead movements during the off-phase indicate that convection flows in the PDMS oil after positioning the bead in front of the needle have not yet come to a complete stop. If this was the case, the measurement was discarded. From the speed of bead movements during the on-phase, the force acting on the particle was then calculated according to Stoke’s law:

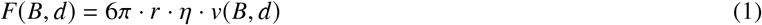

where *r* is the radius of the bead, *η* is the viscosity of the oil, and v is the bead velocity for a given bead-needle distance *d* and magnetic induction *B*. The measured force vs. distance relationship (Fig. 3) followed a stretched exponential function with respect to the bead-needle distance *d* (Eq. 2), whereby the prefactor α and exponent β both depend on the magnetic induction *B*

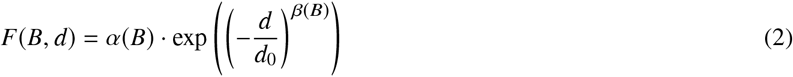

The bead-needle distance *d* in Eq. 2 is given in units of *μ*m, and the normalization of *d* with an arbitrary value of *d*_0_ = 1 *μ*m is introduced for consistency (to remove the physical units).

**Figure 3:**
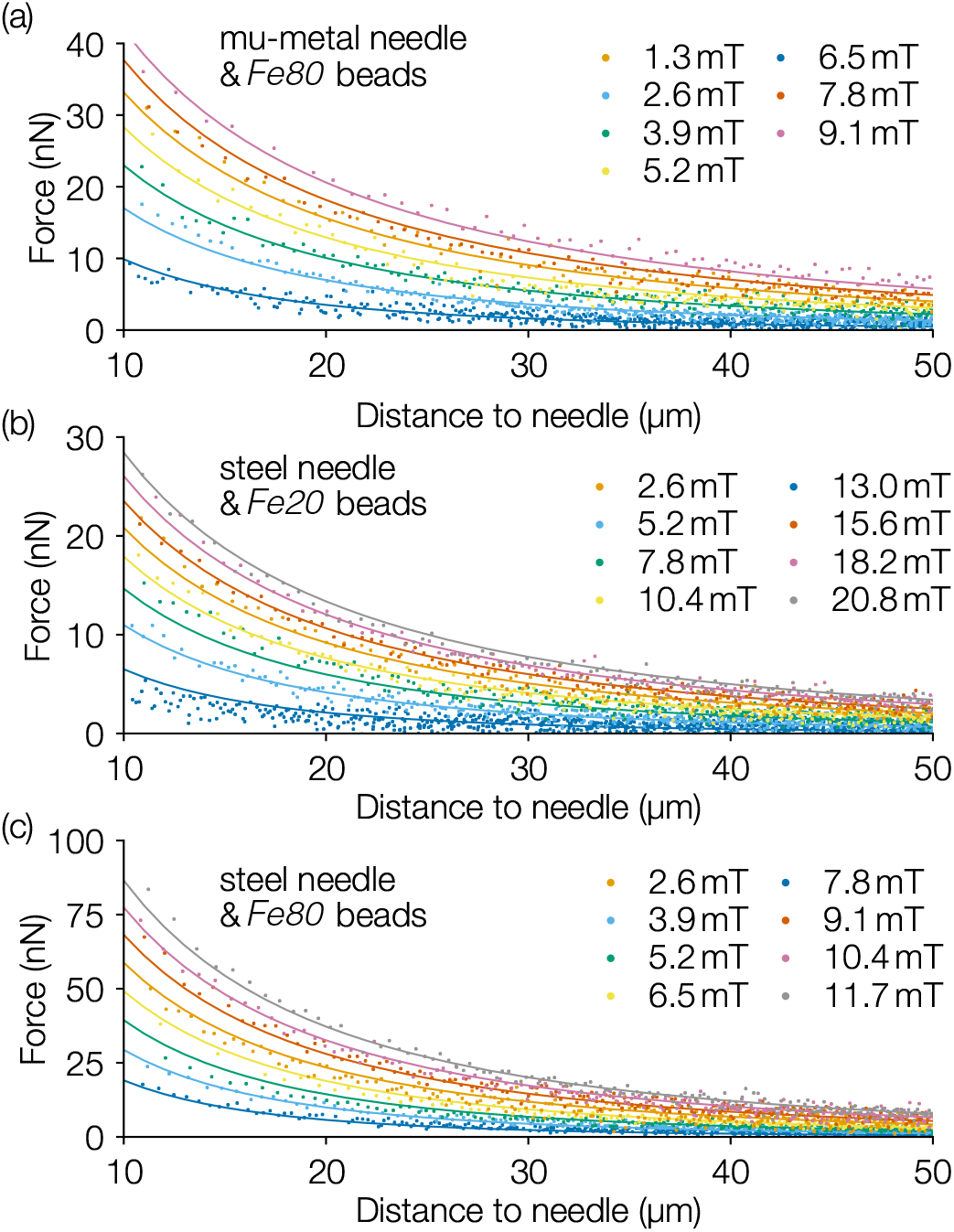
Force calibration. Force-distance curves for different magnetic inductions and for different combinations of tweezer needle material and superparamagnetic beads. (a) Mu-metal needle with carboxylated Fe80 beads, (b) steel needle with epoxylated Fe20 beads, and (c) steel needle with carboxylated Fe80 beads. Solid lines indicate the fits with a stretched exponential function with three parameters (Eq. 2–4).

The prefactor α increased with increasing *B* according to a power-law,

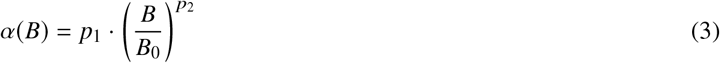

with calibration parameters *p*_1_ and *p*_2_ when the magnetic induction *B* in Eq. 3 is given in units of mT. The normalization of *B* with with an arbitrary value of *B*_0_ = 1 mT in Eq. 3 is introduced for consistency (to remove the physical units).

We also found that the prefactor α to the power of β was constant for all values of the magnetic induction:

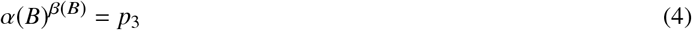

with *p*_3_ being a third calibration parameter. Together, Eq. 2–4 describe the force *F* as a function of magnetic induction *B* and the bead-needle distance *d* with only three fit parameters. We evaluated this simple relationship numerically to compute the magnetic induction *B* necessary to exert a defined force *F* onto a particle located at a given distance *d* from the needle tip. Since the bead-needle distance could be evaluated in real time (from the 50 Hz camera images), and the magnetic induction control loop was sufficiently fast (< 3 ms reaction time), this allowed for real-time force control during experiments.

The values of the fit parameters *p*_1_, *p*_2_, and *p*_3_ depended on the combination of bead type and needle material as shown in Table 1. Accordingly, the combination of Fe80 beads (high iron content and thus high magnetic dipole moment) and St37 core material (high magnetic induction) resulted in the highest forces.

**Table 1:** Calibration parameters for different bead-core material combinations

| core material | bead | $p_1$ | $p_2$ | $p_3$ | force* |
| --- | --- | --- | --- | --- | --- |
| St37 | Fe20 | 78.5 nN | 0.47 | 9.5 | 17 nN |
| Mu-metal | Fe80 | 143.6 nN | 0.52 | 10.0 | 45 nN |
| St37 | Fe80 | 201.8 nN | 0.70 | 17.8 | 75 nN |

\* at 10 $\mu\text{m}$ distance and at a magnetic induction of 10 mT

### Cell culture and sample preparation

Microrheological experiments were conducted using NIH/3T3 murine embryonic fibroblasts. After harvesting with 0.25% trypsin-EDTA (Gibco, Carlsbad, CA), 50,000 cells were seeded in 35 mm tissue culture treated plastic dishes (Nunc, ThermoFisher Scientific, Waltham, MA) and grown overnight in high glucose Dulbecco’s modified Eagle’s medium (DMEM) supplemented with 10% bovine calf serum (BCS, Sigma-Aldrich, St. Louis, MO), penicillin and streptomycin, sodium pyruvate, and L-glutamine. Superparamagnetic Fe80 beads with a diameter of 5.09 *μ*m were coated with the extracellular matrix protein fibronectin (50 *μ*g/ml in PBS overnight at 4 °C) (FN) (21). FN was chosen because of its ability to bind to transmembrane integrin receptors, thus establishing a tight mechanical linkage between the microbead and the cytoskeleton (22). FN-coated beads were sonicated for 15 s, added to the cell culture dish at a 2:1 bead-to-cell ratio, and incubated at 5% CO_2_ at 37°C. After 30 minutes, the cell medium was changed to wash off unbound beads. Measurements were performed at room temperature within a time window of 30 min. Between individual cell measurements, the dish was moved by at least 200 *μ*m to ensure that the cell to be measured had not experienced appreciable forces from preceding measurements.

## RESULTS

### Accuracy analysis

To analyze the accuracy of the magnetic tweezers setup, superparamagnetic beads with a diameter of 5.09 *μ*m (Fe80) were dispersed in PDMS oil with a dynamic viscosity of 28.95 Pa·s and were positioned at 60 *μ*m distance to the de-Gaussed needle tip (St37). We then applied a force protocol consisting of 5 increasing and then decreasing discrete force steps (1, 2, 4, 8, 16, 8, 4, 2, 1 nN), with each step lasting 1 s (see Fig. 4a). From the resulting bead trajectory (blue line in Fig. 4a), the force that acted on the bead was calculated using Stoke’s law (orange points) and compared to the target force (grey area).

**Figure 4:**
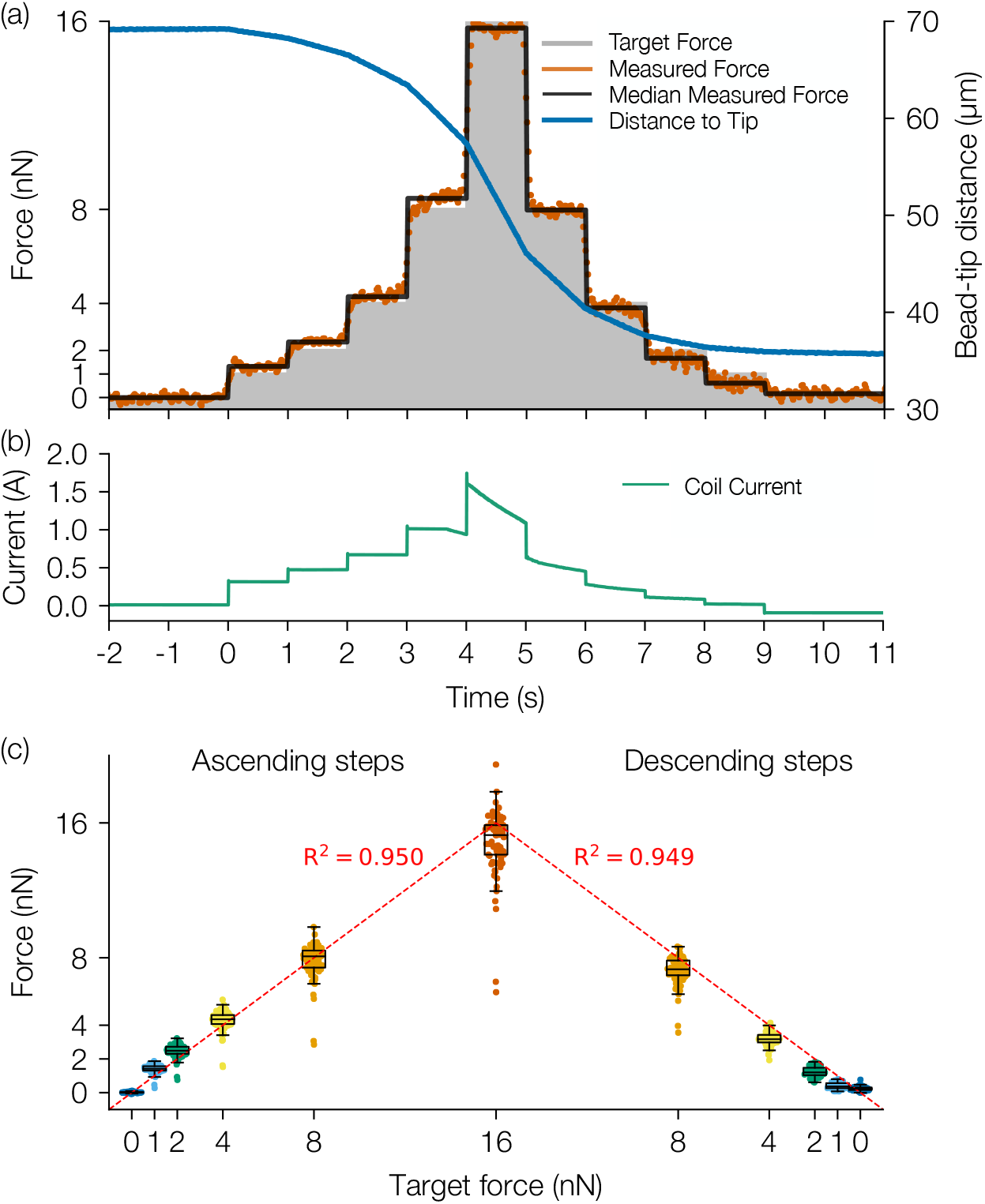
Test with step pyramid protocol. (a) Exemplary bead trajectory (blue) in response to a force step protocol (gray shaded area) for a superparamagnetic bead in PDMS oil (dynamic viscosity of 28.95 Pa·s). The measured force (orange, average shown in black) was calculated from the bead trajectory. (b) Coil current versus time. (c) Measured forces during ascending and descending force steps versus target forces. Points represent forces from individual beads (n = 81), boxplots indicate median values, lower to upper quartile values, and 1.5 interquartile ranges. The line of identity (red dotted line) describes the relationship between measured forces and target forces with a coefficient of determination of *R*^2^ = 0.950 for ascending forces and 0.949 for descending forces.

When averaged over the duration of each force step, the calculated force (black line in Fig. 4a) was close to the target force for both increasing and decreasing steps. The measured coil current (Fig. 4b) increased with higher target forces, as a higher magnetic gradient was necessary to obtain the desired force, but during application of a constant force, the coil current decreased over time as the bead approached the needle tip (blue line in Fig. 4a). Note also that the coil current during the zero force phase prior to the start of the protocol was close to zero, as the needle had been freshly de-Gaussed, but during the zero force phase after completion of the protocol, the coil current was negative to compensate for the remanent magnetic field of the needle (Fig. 4b).

The relationship between measured forces to target forces for ascending force steps was close to the line of identity, with a coefficient of determination of *R*^2^ = 0.950 (Fig. 4c), demonstrating both the quality of the calibration and the quality of the force feedback. Importantly, for descending force steps, we find a similarly high *R*^2^ = 0.949, demonstrating the quality of the hysteresis compensation. The measured force at the end of the protocol (for a zero target force) was slightly increased to 0.23±0.12 nN, n=81 beads, compared to 0.01±0.04 nN at the beginning of the protocol. Note, however, that the beads were considerably closer to the needle tip by the end of the protocol (30±9 *μ*m on average; mean±std) compared to the beginning (61±7 *μ*m on average), and given the exponential relationship between force and bead-tip distance (Eq. 2), even a small remanent field can cause measurable forces. In addition, numerous beads tended to accumulate at the needle tip during force application, which distorts the magnetic field and field gradient. The latter effect should not pose a problem during measurements on cells, as unbound beads that accumulate at the bead tip are washed off prior to starting the experiment.

### Decomposing the compliance of fibroblasts into its viscoeleastic and plastic parts

For cell measurement, the force was stepwise increased every 3 s from 1 nN to 16 nN (with plateau values of 1, 2, 4, 8, and 16 nN) and then stepwise decreased, resulting in a pyramid-like force profile (grey bars in Fig. 5a). The bead position was monitored for 5 s prior to force application (initial phase) and for 10 s after force application (relaxation phase).

**Figure 5:**
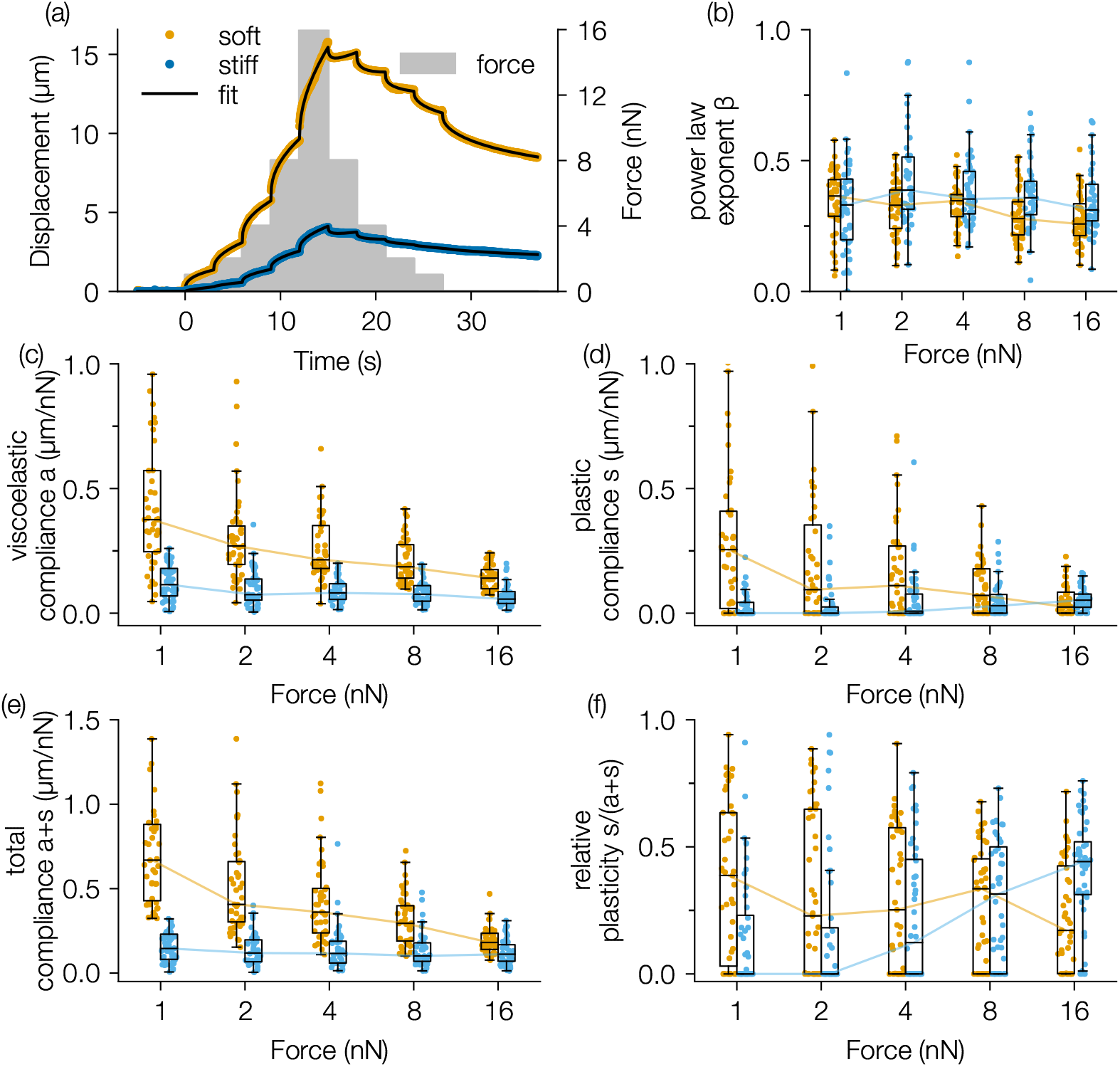
Force-dependency of viscoelastic and plastic cell behavior. (a) Response of a representative cell with high compliance (“soft”: orange dots) and a cell with low compliance (“stiff”: blue dots) to a pyramid of force steps (gray shaded area, 1, 2, 4, 8, 16 nN) fitted with Eq. 5 (black lines). (b) power-law exponent *β*, (c) absolute viscoelastic compliance *a*, (d) absolute plastic compliance *s*, (e) total compliance *a* + *s*, and (f) relative plastic compliance s/(a+s). Data (median, 25%/75% percentiles, 1.5 interquantile range) from soft (orange bars/lines, n = 43 cells) and stiff (blue bars/lines, n = 44 cells) fibroblasts are shown as a function of force.

The deformation of living cells in response to a step-like force typically follows a power-law in time (23, 24). After force removal, the cell returns towards its undeformed shape, also following a power-law in time (24), but the shape recovery is typically incomplete due to irreversible plastic deformations (21). The cell displacement d(t) in response to a single force-on step can be described by a power-law with exponent *β* and a pre-factor (*a* + *s*) that is the sum of the viscoelastic compliance *a* and the plastic compliance *s* (upper part of Eq. 5):

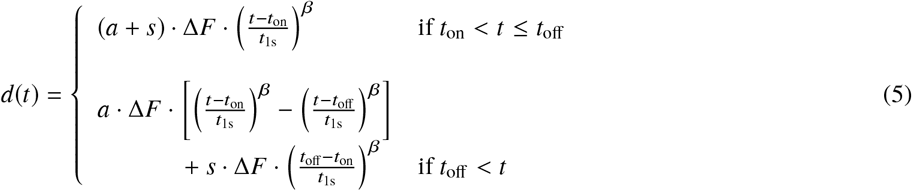

Here, (*t* – *t*_on_) is the time since the start of the force application at *t* = *t*_on_, *t*_1s_ is a consistency factor of 1 s (an arbitrary choice), *t*_off_ is the time at which the force application is turned off, and *F* is the amplitude of the force step. *a* + *s* represents the total compliance, which corresponds to the displacement of a bead after one second of force application, normalized to the applied force.

Upon force removal, the viscoelastic part of the cell’s deformation is reversible and returns gradually to zero (first part of the lower part of Eq. 5), while the plastic part remains constant (second part of the lower part of Eq. 5).

A step pyramid of forces can be described by the superposition of multiple force steps, and the resulting bead displacement is therefore the superposition of bead displacements in response to each individual force step (21, 24) (see Supporting Material S2 for details on the mathematical description). For increasing forces, both the viscoelastic and plastic components of Eq. 5 contribute to the bead displacement towards the needle tip, whereas for decreasing forces, only the viscoelastic component contributes to the bead recoil away from the needle tip while the bead displacements resulting from the plastic components of past force steps are frozen in time.

### Relative plasticity of fibroblasts depends on their initial stiffness

In previous studies, protocols with increasing force steps were used to investigate the force-dependent stiffening of cells (3, 25). These studies demonstrated a stiffening of adherent cells that was proportional to the sum of contractile pre-stress and external stress from the magnetic beads. Accordingly, cells with a high contractile pre-stress, which were stiff at low forces, showed less relative stiffening with increasing forces compared to soft cells. Because protocols with decreasing force steps have thus far not been possible due to the effect of magnetic hysteresis, it is unknown how cell plasticity changes with force, and it is also unknown if the plasticity of soft and stiff cells responds differently to force.

To address these questions, we followed the approach in (25) and grouped cells in a stiff and a soft cohort, with the median of the total compliance at a force of 1 nN as the dividing line (Fig. S3.1a). Only fibroblasts that showed a high coefficient of determination (*R*^2^ > 0.991) and a low error of the fit (square root of the 98% percentile of the squared error < 0.23 *μ*m) were included in the subsequent analysis. All measured data (both included and excluded) and their respective fits are shown in Supporting Material Fig. S4.1–S4.3.

As we measured the bead trajectory in response to an increasing and decreasing force for each individual force step (Fig. 5a), we could fit the viscoelastic (Fig. 5c) and the plastic compliance (Fig. 5d) alongside the power-law exponent *β* (Fig. 5b) using Eq. 5. While the power-law exponent *β* (Fig. 5b) remained approximately constant at around 0.34±0.01 (mean±sem) for all forces for both soft and stiff cells, the viscoelastic and plastic compliance decreased with force for soft fibroblasts (from a=0.38±0.01 *μ*m/nN and s=0.25±0.01 *μ*m/nN at 1 nN to a=0.14±0.01 *μ*m/nN and s=0.02±0.002 *μ*m/nN at 16 nN). Stiff fibroblasts, by contrast, showed a slight decrease in viscoelastic compliance (from 0.11±0.01 *μ*m/nN at 1 nN to 0.06±0.002 *μ*m/nN at 16 nN) and an increase in plastic compliance (from 0.00±0.001 *μ*m/nN at 1 nN to 0.05±0.003 *μ*m/nN at 16 nN).

The total compliance of the stiff cohort was 0.11±0.01 *μ*m/nN at a force of 1 nN and remained nearly constant with increasing force amplitude (Fig. 5e). By contrast, the total compliance of the soft cohort decreased from 0.67±0.02 *μ*m/nN at a force of 1 nN to 0.18±0.01 *μ*m/nN at a force of 16 nN (median±standard error of median). This behavior of pronounced stress stiffening in soft cells compared to more constant responses in stiff cells is consistent with previous findings (25).

The relative plasticity (the ratio *s*/(*a* + *s*) of soft cells showed a similar trend and decreased with increasing force (Fig. 5f), from 0.38±0.02 at 1 nN to 0.17±0.01 at 16 nN. By contrast, the relative plasticity of stiff cells strongly increased with force, from 0.00±0.01 at 1 nN to 0.44±0.002 at 16 nN. This indicates that stiff cells are protected from plastic deformation at low forces, and that they display plastic deformations only at higher forces, whereas in softer cells, large plastic deformations occur already at relatively low forces.

## DISCUSSION

In this study, we present a hysteresis-free magnetic tweezers setup for applying arbitrary force protocols with a maximum force of up to 100 nN. The magnetic induction of the tweezer core is measured with a Hall probe and is feedback controlled so that the magnetic hysteresis of the tweezer core material is effectively eliminated. It is therefore no longer necessary to de-Gauss the tweezer needle between measurements. The relationship between force, magnetic induction and bead-tip-distance can be captured with an analytical equation with only 3 parameters, in contrast to 5 fit parameters that are necessary to capture the relationship for magnetic tweezers with current feedback (11).

The precision with which forces can be applied is predominantly determined by the variability in the iron oxide content between individual beads. During test measurements shown in Fig. 4c for ascending and descending forces, we find that beads are either consistently stronger or consistently weaker at all target forces. The coefficient of variation is 14% at the highest force level of 16 nN, which is similar to the value of 18–28% reported in (11) and which cannot be appreciably reduced further with higher calibration accuracy or better force feedback unless beads with a more uniform iron oxide content are used.

Most importantly, hysteresis-free magnetic tweezers allows for the application of arbitrary force protocols, including protocols with descending forces. Moreover, it is no longer necessary to choose a tweezer core material with low magnetic hysteresis such as Mu-metal alloys that may lose their high permeability when machined or polished, which is frequently required to sharpen the tweezer tip.

We demonstrated the function and versatility of our system by applying a force protocol in the form of a step pyramid to superparamagnetic beads bound to fibroblasts. Such a force step pyramid with ascending and descending steps is suitable for measuring the force-dependence of the total cell compliance (inverse of cell stiffness). In agreement with previous reports (25), we find that soft but not stiff cells become stiffer with increasing external forces. This finding is consistent with the idea that cells are a stress-stiffening material, whereby the total stress is the sum of the internal contractile stress within the cytoskeleton (the so-called pre-stress) and the external stress applied via magnetic beads. Stiff cells have a high pre-stress, and the externally applied magnetic force increases the total stress by only a small fraction - hence these cells do not stiffen further (25, 26). By contrast, in soft cells, the total stress within the cytoskeleton is dominated by the externally applied magnetic forces, and hence soft cells show pronounced stress stiffening.

With a step pyramid-like force protocol of increasing and then decreasing force steps, we can decompose the total compliance into its viscoelastic and plastic components. Similar to the total compliance, we find that the viscoelastic compliance of the softer cell cohort decreases with force, while the viscoelastic compliance remains approximately constant for all forces in the stiff cohort. By contrast, the plasticity of stiff cells tends towards zero for low force and increases with force. In soft cells, the plasticity starts from high values at low forces and steadily decreases with force.

Cell plasticity has been previously shown to originate from bond ruptures within the cytoskeleton, and it can therefore be expected that plasticity increases at higher force, as seen in stiff cells (21). Our data are in agreement with this interpretation. Our observation that the plasticity of soft cells decreases with force suggests that substantial bond rupture is already occurring at low forces, and that more stable cytoskeletal structures such as intermediate filaments prevent further rupturing and yielding events at higher forces. This idea is also supported by our measurements of the power-law exponent of the creep modulus, which reflects the dissipation of elastic energy e.g. from cytoskeletal bond rupture (23). We find that the power-law exponent is only slightly increased in softer cells compared to stiffer cells even at the highest force of 16 nN.

## CONCLUSION

In summary, our results demonstrate the advantages of hysteresis-free magnetic tweezers for studies of cell mechanics, in that it eliminates the need to repeatedly de-Gauss the tweezer needle between measurements, simplifies the force calibration, and allows for the application of arbitrary force protocols such as a pyramid-like ascending and descending step forces.

## AUTHOR CONTRIBUTIONS

**Delf Kah:** Investigation, Visualization, Writing - original draft preparation, Writing - review & editing. **Christopher Dürrbeck:** Methodology, Software, Investigation, Visualization, Writing - original draft. **Werner Schneider:** Methodology. **Ben Fabry:** Project administration, Conceptualization, Methodology, Writing - original draft preparation, Writing - review & editing, Supervision. **Richard Gerum:** Conceptualization, Formal analysis, Visualization, Software, Writing - original draft preparation, Writing - review & editing.

## ACKNOWLEDGMENTS

This work was supported by the National Institutes of Health (HL120839) and the German Science Foundation (DFG FA336/12-1).

## SUPPLEMENTARY MATERIAL

A supplement to this article is included in the LaTeX file and as a separate PDF document.

